# Functions of the Bloom Syndrome Helicase N-terminal Intrinsically Disordered Region

**DOI:** 10.1101/2024.04.12.589165

**Authors:** Colleen C. Bereda, Evan B. Dewey, Mohamed A. Nasr, Jeff Sekelsky

## Abstract

Bloom Syndrome helicase (Blm) is a RecQ family helicase involved in DNA repair, cell-cycle progression, and development. Pathogenic variants in human *BLM* cause the autosomal recessive disorder Bloom Syndrome, characterized by predisposition to numerous types of cancer. Prior studies of *Drosophila Blm* mutants lacking helicase activity or protein have shown sensitivity to DNA damaging agents, defects in repairing DNA double-strand breaks (DSBs), female sterility, and improper segregation of chromosomes in meiosis. Blm orthologs have a well conserved and highly structured RecQ helicase domain, but more than half of the protein, particularly in the N-terminus, is predicted to be unstructured. Because this region is poorly conserved across multicellular organisms, we compared closely related species to identify regions of conservation, potentially indicating important functions. We deleted two of these *Drosophila*-conserved regions in *D. melanogaster* using CRISPR/Cas9 gene editing and assessed the effects on different Blm functions. Each deletion had distinct effects on different Blm activities. Deletion of either conserved region 1 (CR1) or conserved region 2 (CR2) compromised DSB repair through synthesis-dependent strand annealing and resulted in increased mitotic crossovers. In contrast, CR2 is critical for embryonic development but CR1 is not as important. CR1 deletion allows for proficient meiotic chromosome segregation but does lead to defects in meiotic crossover designation and patterning. Finally, deletion of CR2 does not lead to significant meiotic defects, indicating that while each region has overlapping functions, there are discreet roles facilitated by each. These results provide novel insights into functions of the N-terminal disordered region of Blm.

## Introduction

Bloom syndrome helicase (Blm in *Drosophila*; BLM in humans) is an ATP-dependent, RecQ family helicase (5-7). It is conserved across protists, plants, fungi, and animals, with roles in homology-directed DNA repair (HDR), cell-cycle progression, meiosis, and development (1, 3, 8-16). Pathogenic variants in *BLM* cause Bloom Syndrome, a rare autosomal recessive disorder characterized by a high predisposition to a broad range of cancers, sun sensitivity, short-stature, sterility, and immunodeficiency (5, 17, 18). *BLM* mutations have also been found in sporadic cancers (19-22). The high predisposition to cancer in individuals with Bloom Syndrome is associated with genome instability, including high rates of exchange between sister chromatids and homologous chromosomes (23, 24).

One important function of BLM/Blm in HDR is disassembly of DNA repair intermediates, which is done in concert with topoisomerase III alpha (TopIIIα) (25-28). BLM and TopIIIα, together with RMI1 (which Drosophila lacks (29)), unwind D-loops to promote dissociation of the invading strand in synthesis-dependent strand annealing (SDSA) (2, 7, 25, 27) and catalyzes dissolution of double Holliday junctions (dHJs) (13, 26, 30-32). These two functions prevent mitotic crossovers and therefore minimize loss of heterozygosity (LOH) and chromosome rearrangement. Blm orthologs also have functions in meiosis, but these include promoting crossovers (reviewed in 14). In *Drosophila*, loss of Blm results in decreased meiotic crossover rates, compromised crossover distribution and increased chromosome mis-segregation (non-disjunction) (1).

BLM/Blm also has functions in repair of stalled replication forks to promote an efficient S-phase (12, 25). BLM accumulates at stalled forks along with other DNA repair regulators, with *in vitro* studies suggesting BLM may act to regress stalled forks behind a DNA lesion to promote lesion removal by other repair pathways (33, 34). A second BLM cell cycle role is to resolve anaphase bridges to allow proper chromosome segregation during mitosis (35, 36). In human cells, this activity is mediated through interaction with topoisomerase IIα (TopIIα) (37). Micronuclei and aneuploidy are more prevalent in *BLM*-deficient cells, underscoring the importance of this BLM role to genome stability (38, 39). In *Drosophila*, embryos lacking Blm have increased anaphase bridges during rapid syncytial cell cycles, resulting in high rates of embryonic death (3, 16). These various functions suggest that BLM/Blm regulation is dependent on cell type and developmental timing.

While BLM/Blm is best known by its RecQ helicase domain, there are large, intrinsically disordered regions (IDRs) both N- and C-terminal to this domain (Figure 1A). These regions, though poorly conserved in primary sequence, are likely candidates for both regulatory modifications and protein-protein interactions. TopIIIα is thought to bind in at least one of these regions, and other proteins’ interactions have been mapped to them as well (28, 37, 40). Despite this, the IDRs have been relatively poorly explored compared to the helicase domain, even though they make up more than half of the protein (Figure 1A). A study in *Drosophila* underscores the importance of these regions, with a *Blm* allele that deletes most of the unstructured N-terminus (*Blm*^*N2*^) compromising HDR and meiotic roles while only mildly affecting early embryonic functions (Figure 1B) (3).

**Figure 1.**
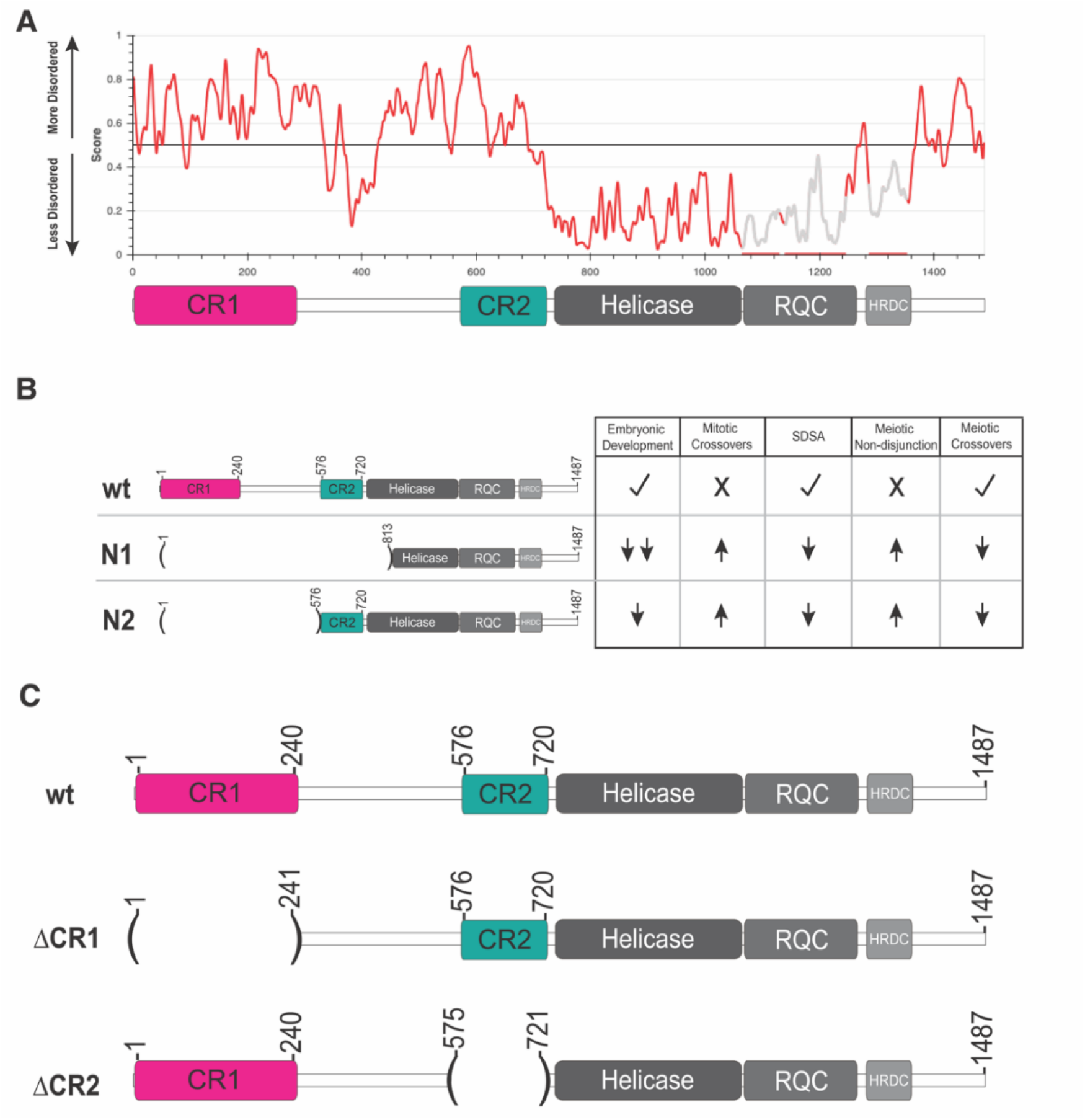
Bloom syndrome helicase (Blm) predicted structural order and alleles used. **(A)** IUPred3 (2) plot predicting the ordered and intrinsically disordered regions of Blm. A schematic of Blm domains is placed below for reference, illustrating that conserved regions 1 and 2 (CR1 and CR2) are predicted to be mostly disordered (>0.5). **(B)** Previously characterized alleles of *Blm* and their effects on Blm functions relative to wild-type (wt). A null allele, *Blm*^*N1*^ (N1) eliminates all well-characterized Blm functions, while the separation-of-function allele *Blm*^*N2*^ (N2) only moderately affects embryonic development. **(C)** Schematic of Blm deletions used compared to the wt Blm protein. ΔCR1 deletes amino acids 2-240, preserving amino acids 241-1487 in frame; ΔCR2 deletes amino acids 576-720, fusing amino acids 1-575 and 721-1487 together in frame.

To further investigate the function of the intrinsically disordered N-terminal region, we characterized the impacts of deletions of two N-terminal regions conserved in closely related *Drosophila* species on embryonic development, HDR, and meiosis. We find that while deletion of the first 240 amino acids does not compromise Blm in meiotic chromosome segregation, it does affect embryonic development, HDR, and meiotic crossover distribution, albeit less severely than *Blm* null mutations. A deletion of the 146 amino acids just prior to the start of the structured RecQ helicase domain results in severe defects in cell division and development but has milder effects on HDR and apparently normal meiotic crossover distribution and segregation. These findings highlight the importance of investigating intrinsically disordered Blm regions to understanding function.

## Results

### Identification of N-terminal regions conserved among *Drosophila* species and deletion by CRISPR/Cas9 genome editing

Despite high conservation in the helicase domain of Blm, the roughly 720 amino acid N-terminal region is not well conserved among multicellular organisms. This region is predicted to be intrinsically disordered (Figure 1A). Prior studies the *Blm*^*N2*^ allele, which deletes the first 575 residues of the IDR but retains 146 residues upstream of the helicase domain pointed to a potential role of this helicase-adjacent N-terminal region in embryonic development (McVey, 2007). To further examine functions of the N-terminal region, we narrowed our focus to conservation among more closely related *Drosophila* species (Figure S1). Alignment of these species identified two regions of high similarity, which we term conserved region one (Figure 1C, CR1; amino acids 1-240) and conserved region two (Figure 1C; CR2; amino acids 533-720). CR1 may contain one of the two regions in human BLM found to interact with TopIIIα (28). We further narrowed CR2 to contain only the N-terminal amino acids predicted to present in the protein produced by the *Blm*^*N2*^ allele (amino acids 576-720), to compare their functions more directly. Using CRISPR/Cas9 genome editing, we separately deleted the sequences encoding amino acids 1-240 and 576-720 in the endogenous *Blm* gene (Figure S2). We refer to these alleles as *Blm*^Δ*CR1*^ and *Blm*^Δ*CR2*^.

### Embryonic hatch rates are affected differently by each N-terminal Blm deletion

The absence of maternally supplied Blm results in frequent anaphase bridges and high rates of embryonic lethality (3, 16). To determine the effects of each deletion on Blm function in embryonic development, we conducted embryonic hatching assays. In agreement with prior results, embryos from females homozygous for the *Blm*^*N1*^ allele, which does not produce Blm transcript or protein (3), have severely reduced hatch rates. In contrast, there is much smaller, though significant, reduction in hatching of embryos from *Blm*^*N2*^ mothers (Figure 2). Functionality of the Blm^N2^ protein in embryogenesis likely requires the presence of the helicase, RecQ, and HRDC domains, but the predicted Blm^N2^ protein also has the last 146 residues of the N-terminal IDR that may contribute to function. We assessed the effects of deletion of this region (CR2) on embryogenesis (Figure 2). Strikingly, embryos from *Blm*^Δ*CR2*^ females have a low hatch rate similar to that of embryos from *Blm*^*N1*^ mothers, consistent with this region being critical to Blm function in embryonic development.

**Figure 2.**
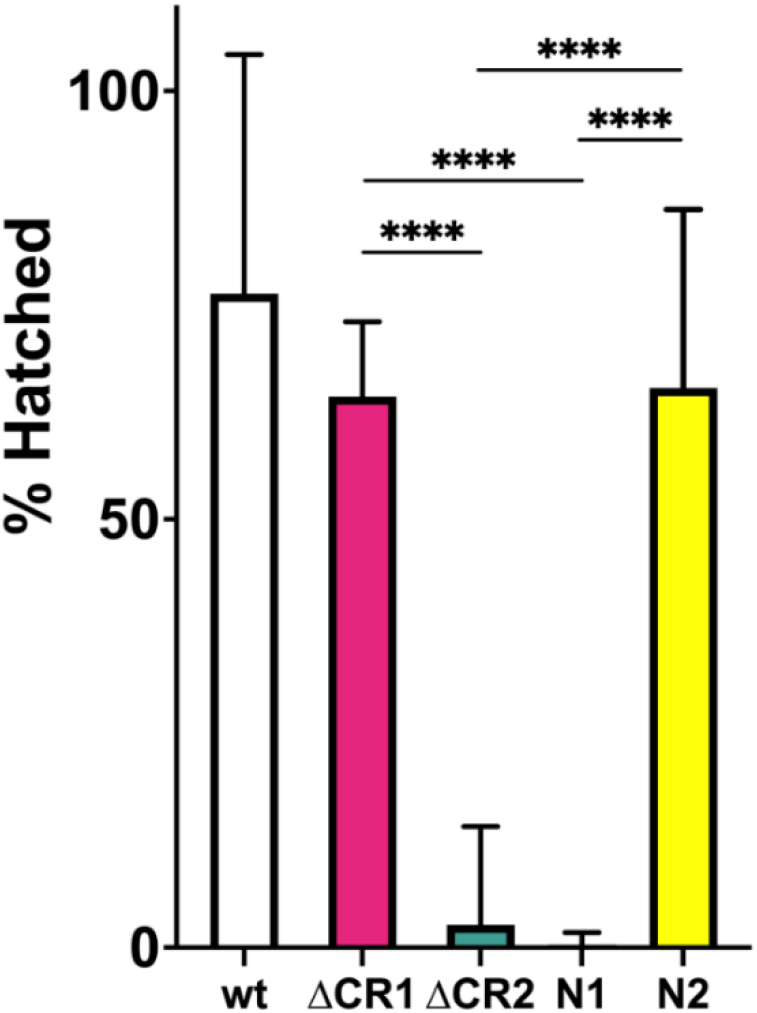
Hatching of embryos from *Blm* mutant mothers. Virgin females homozygous for the *Blm* alleles indicated on the X-axis were crossed to Oregon-RM males and allowed to lay overnight on grape-juice agar. Embryos were transferred to fresh grape-juice agar plates and scored for hatching 48 hours later. Each experiment was repeated three times, with 100-250 embryos transferred each time. Embryos from *Blm*^Δ*CR2*^ (ΔCR2) or *Blm*^*N1*^ (N1) females are rarely able to complete development. Embryos from *Blm*^Δ*CR1*^ (ΔCR1) and *Blm*^*N2*^ (N2) have a modest but significant reduction in hatch rates. We conclude that the CR2 region is more critical for embryonic development but the CR1 region contributes only to a small degree. *n* = wt: 598; ΔCR1: 1080; ΔCR2: 743; N1: 706; N2: 700. **** *p* < 0.0001 by Fisher’s exact test.

We also assayed the effects of the CR1 deletion on hatching. While the fraction of embryos from *Blm*^Δ*CR1*^ females that hatched was significantly lower that of embryos from wild-type females, it was significantly higher than that of embryos from either *Blm*^Δ*CR2*^ or *Blm*^*N1*^ females (Figure 2), indicating that this region is less important to Blm roles in embryonic development. This was consistent with the high hatch rate of embryos from *Blm*^*N2*^ mothers, which also significantly lower than that of wild-type but higher than that of *Blm*^Δ*CR2*^ and *Blm*^*N1*^, in line with previous findings (CITE McVey 2007).

### Mitotic crossovers are moderately elevated in *Blm*^Δ*CR1*^ and *Blm*^Δ*CR2*^ mutants

Flies with the *Blm*^*N1*^ or *Blm*^*N2*^ deletion have elevated spontaneous mitotic crossovers, probably due at least in part to compromised SDSA and/or dHJ dissolution functions (Figure 1B) (3, 11). We assayed *Blm*^Δ*CR1*^ and *Blm*^Δ*CR2*^ mutants and found they also have elevated mitotic crossovers (Figure 3), but at rates (0.28% and 0.61%, respectively) that are significantly lower than those of *Blm*^*N1*^ and *Blm*^*N2*^ alleles (2.3% and 2.4%, respectively). This suggests that loss of CR1 or CR2 allows some non-crossover repair or some other function that prevents lesions that can be repaired as crossovers.

**Figure 3.**
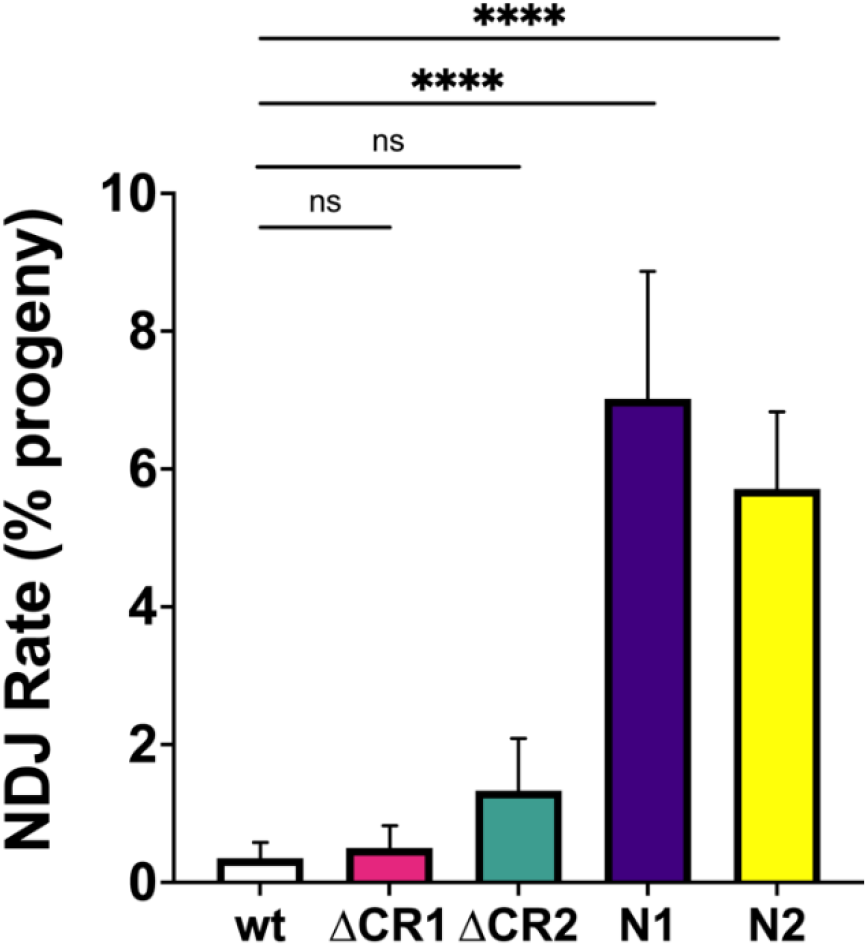
Meiotic non-disjunction (NDJ). Virgin females with the *Blm* alleles indicated on the X-axis over the *Blm*^*N1*^ null allele (*Blm*^*N1*^ was over *Blm*^*D2*^) were crossed to *y sc cv v g f / Dp(1;Y)B*^*S*^ males in at least 15 vials, each serving as a biological replicate. Progeny were scored for non-disjunction (NDJ), indicated by bar eyes in daughters (*XXY*) and non-bar-eyes in sons (*X0*) genotypes. The number of NDJ progeny was doubled to correct for genotypes that do not progress to adulthood (*XXX* and *Y0*), then NDJ rate was determined as a ratio of the number of corrected NDJ individuals to total progeny for each genotype. Neither *Blm*^Δ*CR1*^ (ΔCR1) nor *Blm*^Δ*CR2*^ (ΔCR2) had a significant increase in NDJ compared to wild type (wt). In agreement with prior studied, both *Blm*^*N1*^ and *Blm*^*N2*^ females have significantly elevated NDJ. Number of progeny = wt: 6900; ΔCR1: 3593; ΔCR2: 777; N1: 2959; N2: 1906. **** *p* < 0.0001; ns: *p* > 0.05 by the methods described in Zeng *et al*. (4).

### CR1 and CR2 are required for repair of DSBs by SDSA

Blm has a key role in SDSA, where it is thought to promote dissociation of D-loops during or after synthesis (7, 41). To determine whether the lower number of mitotic crossovers in the *Blm*^Δ*CR1*^ and *Blm*^Δ*CR2*^ mutants relative to null mutants is due to better capabilities of these alleles to complete SDSA, we conducted the *P*{*w*^*a*^} SDSA assay (7, 41). In this assay, effectiveness of SDSA in the male germline is determined by scoring progeny for a red eye color that indicates synthesis of >4000 bp from each end of gap generated by transposase-mediated excision, followed by dissociation of nascent strands and annealing of an internal repeat (the long terminal repeat of a *copia* retrotransposon). This outcome is greatly reduced in *Blm*^*N1*^ and *Blm*^*N2*^ mutants, demonstrating inability to complete SDSA (3, 7). We found a similar reduction in *Blm*^Δ*CR1*^ and *Blm*^Δ*CR2*^, (Figure 4), revealing a requirement for both CR1 and CR2 in SDSA repair.

**Figure 4.**
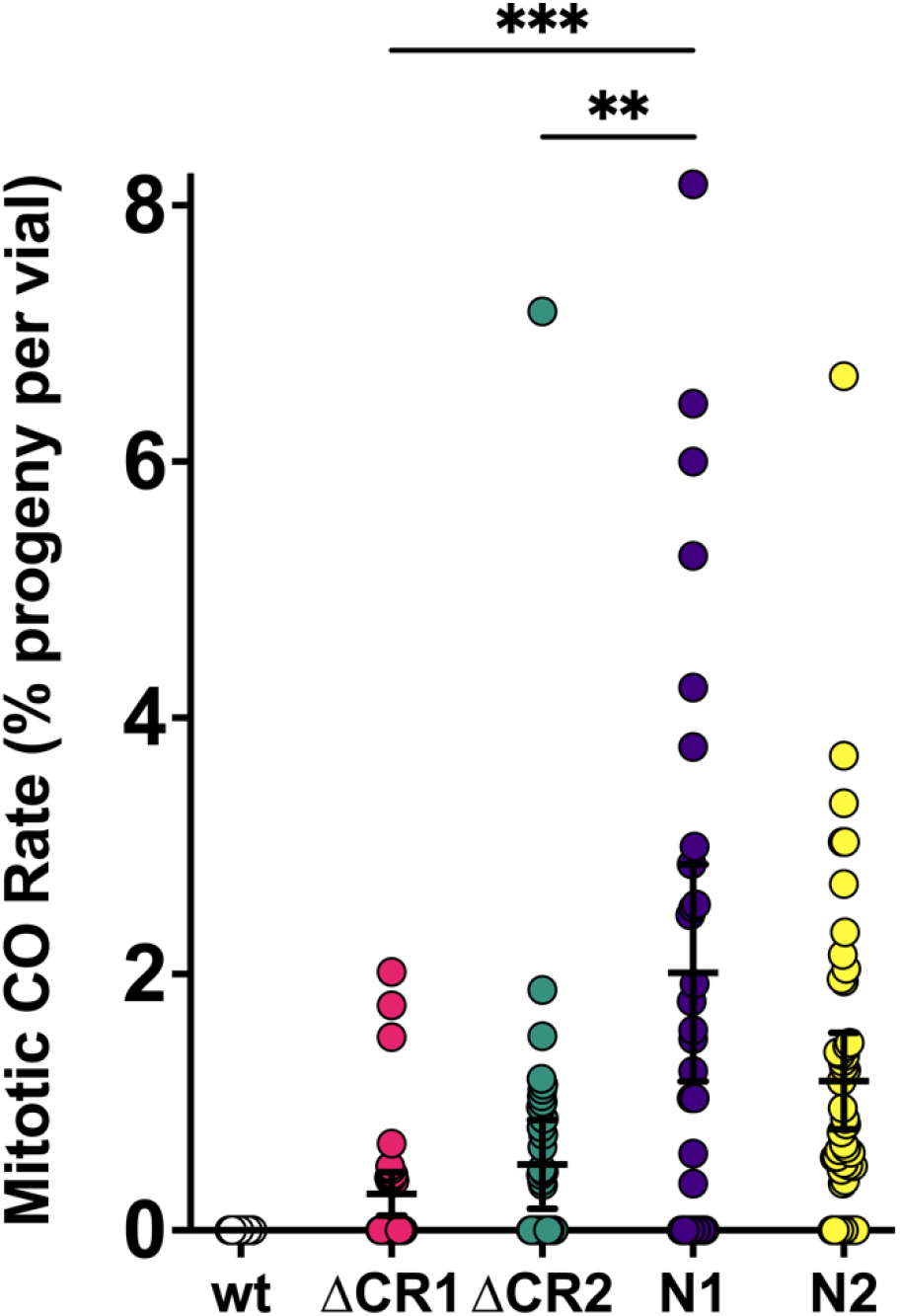
Mitotic crossovers in *Blm* mutants. Single males with the *Blm* alleles indicated on the X-axis over a the *Blm*^*N1*^ null allele (or *Blm*^*D2*^ for *Blm*^*N1*^) were crossed to homozygous *net dpp*^*ho*^ *dpy b pr cn* recessive phenotypic marker virgin females, with each vials serving as a biological replicate. Progeny were then scored for mitotic crossovers occurring in the parental male’s germline, indicated by mixed presence and/or absence of recessive phenotypes. To obtain the mitotic crossover rate per vial, the number of mitotic crossover progeny was divided by the total number of progeny in that vial. Rates for each vial were then pooled to obtain a mean mitotic crossover rate for each genotype. Crossovers are extremely rare in wild-type males (3), so these are excluded from statistical analyses. While both *Blm*^Δ*CR1*^ (ΔCR1) and *Blm*^Δ*CR2*^ (ΔCR2) have mitotic crossovers, the rates in both mutants are significantly less than that of the *Blm*^*N1*^ null mutants (N1). \*\*\**p*<0.001 and \*\**p* < 0.01 by ANOVA with Tukey’s Post Hoc. Compared to the separation-of-function *Blm*^*N2*^ (N2; *n* = 7390) mutant, ΔCR1 mutants had significantly fewer mitotic crossovers (*p* < 0.05 by ANOVA with Tukey’s Post Hoc), but ΔCR2 was not significantly different. *n* = wt: 37 vials, 7091 progeny; ΔCR1: 54 vials, 9284 progeny; ΔCR2: 44 vials, 7174 progeny; N1: 88 vials, 9368 progeny; N2: 54 vials, 7390 progeny.

### *Blm*^Δ*CR1*^ and *Blm*^Δ*CR2*^ mutants have distinct meiotic phenotypes compared to *Blm*^*N1*^ null mutants

Loss of Blm causes meiotic non-disjunction (NDJ; Figure 1B) (1, 3). To assess this function in our *Blm* deletion alleles, we performed an *X* chromosome NDJ assay. The rates of NDJ in *Blm*^Δ*CR1*^ (0.5%) and *Blm*^Δ*CR2*^ females (1.33%) were not significantly different from that of wild-type females (Figure 5), indicating that the regions deleted in CR1 and CR2 are dispensable for Blm functions that prevent NDJ. *Blm*^Δ*CR1*^ and *Blm*^Δ*CR2*^ each also had significantly lower NDJ rates than *Blm*^*N1*^ and *Blm*^*N2*^ females (7.02% and 5.71%, respectively).

**Figure 5.**
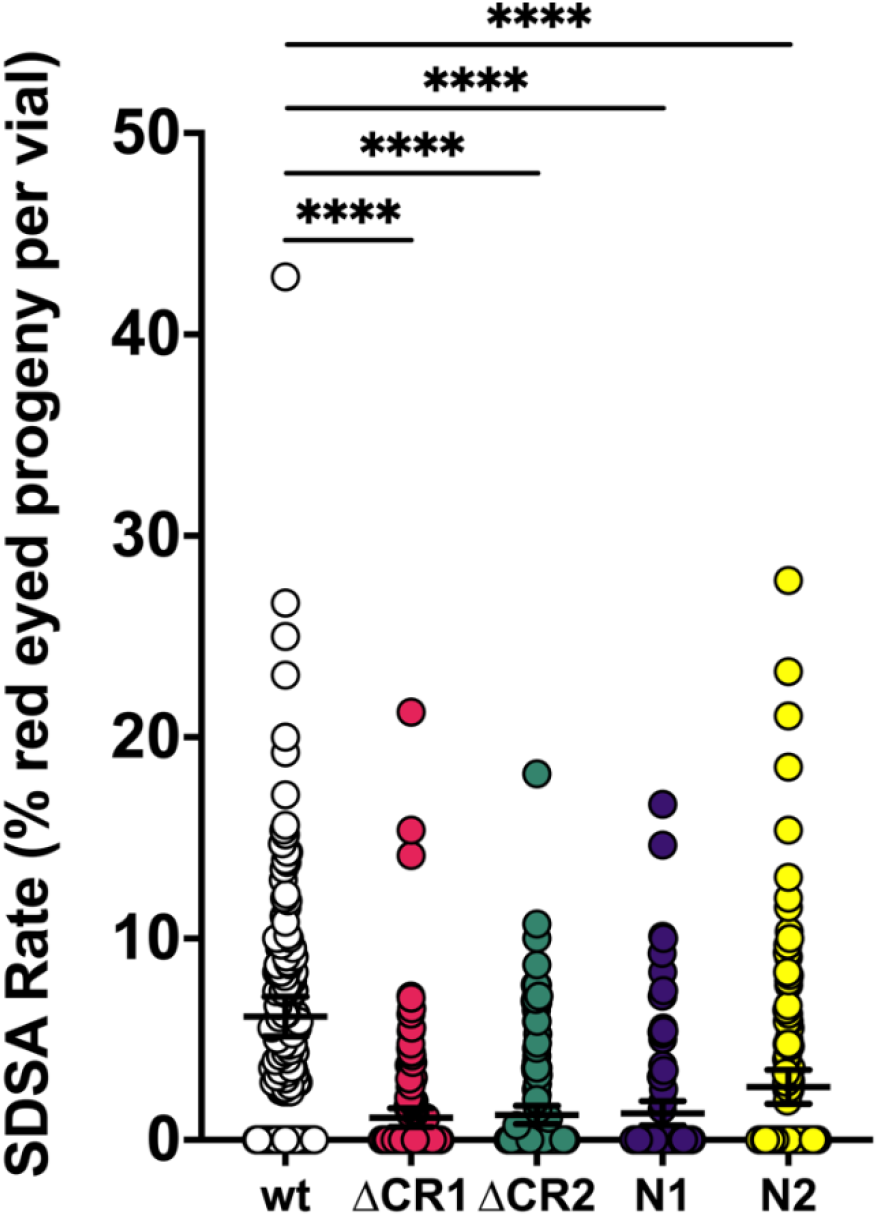
Repair of DNA gaps by SDSA. Single males with the *Blm* alleles indicated on the X-axis (in *trans* to a *Blm*^*D2*^ null allele) and the Δ2-3 transposase were crossed to homozygous *P*{*w*^*a*^} virgin females, with each vial serving as a biological replicate. Progeny without the Δ2-3 transposase were scored for the type of repair that occurred in the parental male’s germline, with red eyes indicating completed SDSA, yellow or white eyes indicating end-joining, and apricot eyes indicating either no excision or repair that restored the complete *P*{*w*^*a*^}. SDSA frequency is the percentage of proteny with red eyes. All mutants had significantly lower numbers of red-eyed progeny than wild-type. \*\*\*\**p*<0.0001 by ANOVA with Tukey’s Post Hoc and Kruskal-Wallis with Dunn’s Multiple comparisons. *n* =wt: 151 vials, 4675 progeny; ΔCR1: 45 vials, 6393 progeny; ΔCR2: 148 vials, 4328 progeny; N1: 106 vials, 4197 progeny; N2: 133 vials, 3860 progeny.

We also wanted to examine crossing over in *Blm*^Δ*CR1*^ and *Blm*^Δ*CR2*^ mutants. Based on results from the NDJ assay, we hypothesized that both *Blm*^Δ*CR1*^ and *Blm*^Δ*CR2*^ mutants would have normal meiotic crossovers. Surprisingly, crossovers were significantly reduced in *Blm*^Δ*CR1*^ mutants (total genetic length of the region assayed was 44.8 cM in *Blm*^Δ*CR1*^ vs. 52.4 cM in wild-type, *p* < 0.0001), particularly in the middle of the chromosome arm assayed (Figure 6). Also surprising was that *Blm*^Δ*CR2*^ mutants had significantly more crossovers (55.3 cM vs. 52.4 cM in wild-type females), with a similar distribution (Figure 6). These results suggest that CR1 and CR2 have different functions in meiosis, contributing to meiotic crossover distribution in opposing ways.

**Figure 6.**
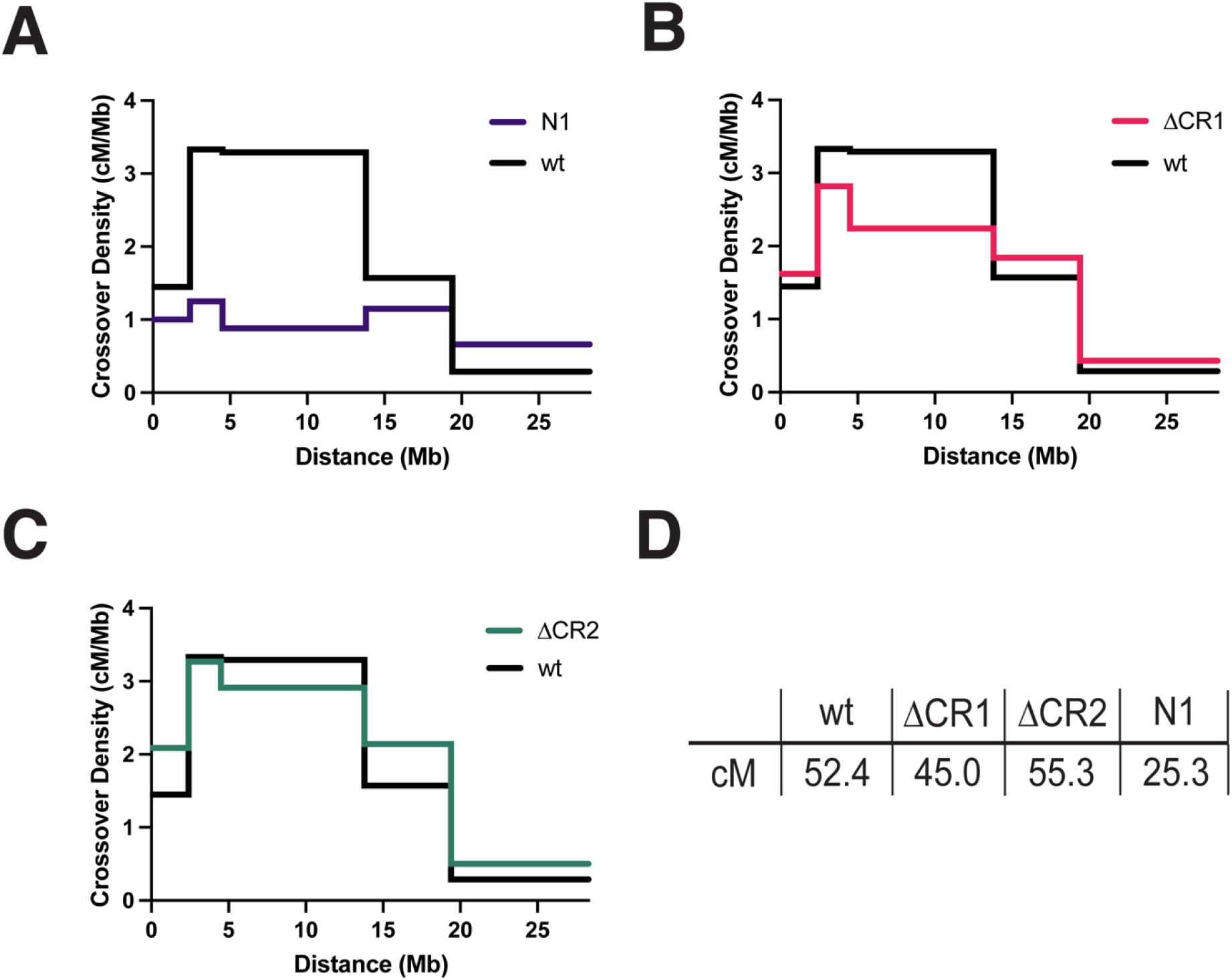
Meiotic Crossovers in *Blm* Mutants. Virgin females with the *Blm* alleles indicated on the X axis (in *trans* so the *Blm*^*N1*^ null allele or, for *Blm*^*N1*^, *Blm*^*D2*^) and heterozygous for the *net dpp*^*ho*^ *dpy b pr cn* chromosome were test crossed and progeny were scored for recessive phenotypes. Graphs show crossover density (cM/Mb) for each genetic interval. **(A)** *Blm*^*N1*^ (N1) had a significant reduction in crossovers and an altered distribution, in agreement with a prior study (1). **(B)** Crossovers were significantly reduced in *Blm*^Δ*CR1*^ (*p* < 0.01 by Fisher’s exact test). **(C)** *Blm*^Δ*CR2*^ mutants had a modest but statistically significant increase in crossovers (*p* < 0.01 by Fisher’s exact test. **(D)** Both *Blm*^Δ*CR1*^ and *Blm*^Δ*CR2*^ mutants had significantly higher crossing over than *Blm*^*N1*^ (*p* < 0.0001 fof each, Fisher’s exact test for each. *n* =4031 for wt; 4049 for *Blm*^Δ*CR1*^; 5088 for *Blm*^Δ*CR2*^.

## Discussion

### CR2 is required for embryonic development

We have shown here that two previously uncharacterized regions of *Drosophila* Blm have distinct functional roles. Embryos from *Blm*^Δ*CR2*^ homozygous mutant females show compromised hatching, to a similar degree as null mutants. This is likely due to the accumulation of anaphase bridges resulting from defects in rapid replication and/or an inability to resolve sister chromatid entanglements during anaphase. Russell and colleagues (37) mapped a TopIIα interaction with human BLM to the region that may correspond to CR2 of *Drosophila* Blm, but this interaction has not been mapped in *Drosophila* Blm.It is possible that this region is regulated to either promote or prevent such interaction. Phosphorylation by ataxia-telengiectasia and *rad3*^*+*^ related and mutated (ATR/ATM) kinases might be one way to promote interaction with TopIIα as part of the DNA damage response, both in stalled fork repair and resolution of anaphase bridges. Human ATR phosphorylates BLM at two residues to promote the recovery of replication forks after stalling by hydroxyurea, and mutation of these residues to alanine results in cell cycle arrest (42). Tangeman and colleagues (43) found that additional predicted ATR/ATM phosphorylation sites are important for BLM nucleolar localization and TopI interaction. The CR2 region has several S/T-Q sites that are possible targets of ATR/ATM phosphorylation, but a *Drosophila* phosphoproteomic analysis did not identify any phosphopeptides from this region in embryos (44).

Harris-Behnfeldt and colleagues (45) showed a potential requirement for phosphorylation of the human BLM region analogous to *Drosophila* Blm CR2, identifying several residues that when mutated to alanine increase ultra-fine anaphase bridges and DNA double-strand breaks while decreasing colocalization of BLM and TopIIα. While some of these residues were predicted to be phosphorylated by ATR/ATM, others were not, suggesting that regulation may be distinct in different species. Regardless of the kinase, regulation and specifically phosphorylation of this region is important to BLM/Blm interaction with TopIIα and function in replication fork repair and resolution of anaphase bridges, even if the residues and kinases involved differ.

### CR1 and CR2 are required for SDSA and prevention of mitotic crossovers

Both *Blm*^Δ*CR1*^ *and Blm*^Δ*CR2*^ mutants showed defects in DSB repair. SDSA rates were compromised to the same extent as in *Blm*^*N1*^ null mutants, but the frequency of spontaneous mitotic crossovers was not as high as in null mutants. One possibility is that CR1 and CR2 are required for SDSA but not for dHJ dissolution. It is not possible to test this possibility *in vivo* due to the lack of a dHJ dissolution assay. CR1 may be analogous to the major TopIIIα-interacting region of human BLM (28). *In vitro*, dHJ dissolution requires TopIIIα, which might suggest that *Blm*^Δ*CR1*^ mutants would be defective for dissolution; however, human TopIIIα also interacts with the C-terminus of BLM (28). Interactions between *Drosophila* Blm and TopIIIα have not been mapped. Furthermore, although BLM can disassemble short D-loops *in vitro*, it is likely that disassembly of D-loops *in vivo*, where the ends are not free to rotate, requires topoisomerase activity, so loss of this interaction may impair both SDSA and dHJ dissolution.

How then might each of the Blm deletions studied lead to compromised SDSA? For CR1, it may be that Blm-TopIIIα interaction with both N- and C-terminal regions of Blm together are necessary for effective SDSA, with loss of either leading to disrupted repair. This could be further explored with a C-terminal deletion in examination of SDSA and mitotic crossovers. We attempted to pursue such a mutant, deleting both the final 100 and 150 amino acids in the unstructured C-terminal region of Blm, but both deletions were homozygous lethal, which is unexpected given that *Blm* null mutants are viable. Future studies could also target ATR/ATM predicted phosphorylation residues within CR1 to attempt to characterize the role of regulation of this region in effective SDSA.

As for the role of CR2 in SDSA, it may be that an interaction with TopII is required for this process. While TopIIIα is likely the primary topoisomerase involved in dissolution of D-loops in SDSA, TopII may be necessary to decatenate more complex DNA structures resulting from errors or disrupted repair. Future directions will also work to characterize the effects of regulation of CR2 on SDSA, with positive regulation potentially promoting additional interaction and/or stabilization.

### CR1 and CR2 contribute to distinct meiotic processes

The two deletions caused different meiotic phenotypes. *Blm*^Δ*CR1*^ mutants had a significant reduction in meiotic COs, whereas *Blm*^Δ*CR2*^ mutants had an increase. Neither mutant had increased NDJ. These are both different from *Blm* null mutants, which have decreased meiotic COs, altered CO distribution, and elevated NDJ.

CR1 appears to play a role in meiotic CO distribution, but in a way that is not required for proper segregation of meiotic chromosomes. How loss of this region impacts crossovers but not segregation is unknown. While many of the components involved in meiotic and mitotic DNA repair are conserved, their regulation does often differ in each process. CR1 would be hypothesized to be involved in the resolution of meiotic DSBs as COs, but not in their repair as NCOs. This would be explained by a higher incidence of meiotic NCOs in *Blm*^Δ*CR1*^ mutants. This might be detectable in whole-genome sequencing of progeny to quantify NCOs. Ability of Blm^ΔCR1^ to resolve any bridged chromosomes during meiotic anaphases could explain the normal NDJ numbers. We should note too that while *Blm*^Δ*CR1*^ CO numbers were significantly lower than wild-type, they were much higher than *Blm* null mutants, so the effects on COs may be mild enough to lead to normal meiotic chromosome segregation.

Blm^ΔCR2^ meiotic activities are also unusual, with a significant increase of meiotic COs yet normal meiotic disjunction. We speculate that this may be due to an inability of Blm^ΔCR2^ to resolve DSBs as NCOs, sending more of them into a CO pathway. This would be consistent with CR2, but not CR1, being required for meiotic SDSA and/or dHJ dissolution. Consistent with this hypothesis, overall numbers and patterning of crossovers would not be disrupted, possibly due to Blm^ΔCR2^ having an intact CR1.

### Conclusion

We have assessed genetic functions of N-terminal, unstructured regions of *Drosophila* Blm helicase. We show that deletion of the first 240 amino acids (CR1) does not impair embryonic development or meiotic chromosome segregation but disrupts mitotic DNA repair and meiotic crossover distribution. Deletion of the 146 amino acids upstream of the helicase domain (CR2) leads to severely disrupted embryonic development and aberrant mitotic DNA repair but allows normal meiotic crossover distribution and chromosome segregation. Through this characterization, we have begun to assign distinct Blm functions to different regions of the N-terminus, leading to a better understanding of how this complex protein works to promote development, meiosis, and genome stability.

## Methods

### CRISPR/Cas9 Deletion of CR1 and CR2

The endogenous CR1 and CR2 region of the *Blm* gene (chromosome *3L*, cytological region 86E17) were deleted in-frame using CRISPR/Cas9 genome engineering similar to that described in Lamb et al., 2017 (Figure S2). A plasmid containing DNA homologous to 5’ and 3’ *Blm* flanking sequence of either CR1 or CR2 (pSL1180 ΔCR1 5’+3’ Homology Arms and pSL1180 ΔCR2 5’+3’ Homology Arms, respectively) and another plasmid containing 5’ and 3’ *Blm* gRNAs for CR1 or CR2 were (pCFD4 *Blm* 1+240 gRNA and pCFD4 *Blm* 576+720 gRNA, respectively) were simultaneously injected into *Drosophila* embryos expressing Cas9 in their germline stem cells under control of the *nanos* promoter (Genetivision, Houston, TX). Upon eclosion of these embryos, single male progeny were crossed to *TM3,Sb/TM6B, Hu Tb* females to balance their potentially edited chromosomes. Once balanced, subsequent single male progeny were again mated to *TM3,Sb/TM6B, Hu Tb* females. After being allowed to mate for 3-4 days at 25 °C, these single males were collected, frozen, and had their genomic DNA isolated to screen for successful deletions by PCR. For vials in which parental males contained the deletion (indicated by a smaller DNA band after PCR compared to wild-type flies), progeny were then mated to siblings to establish a stock. Each deletion stock was then further screened via genomic extraction, PCR, and sequencing of homozygous flies within the resulting stock to confirm the deletion resulted in the correct sequence and that there were no frameshifts. All homozygotes sequenced from each resulting stock contained the correct deletion, flanking sequence, and were not frameshifted, indicating that CR1 and CR2 were successfully deleted.

### Embryonic Hatching Assay

20-30 virgin females homozygous for each *Blm* allele were crossed to 15 *Oregon-RM* (wild-type) males and allowed to acclimate to grape-juice agar plates with yeast paste for 24-36 hrs at 25 °C. Plates were then changed and embyros were collected overnight (16 hrs) at 25 °C. 150-300 embryos were then transferred to a gridded grape juice agar plate (10/grid) and scored for hatching after 48 hours at 25 °C. Hatch assays were completed in three replicates for each allelic condition, with a minimum of 550 total embryos assayed per condition.

### Meiotic Non-disjunction Assay

Female meiotic non-disjunction (NDJ) of the *X* chromosome was measured by first crossing *w; Blm*^*N1*^*/TM3, Sb* virgin females to *Oregon-RM* (wild-type) males or males with the *Blm* allele of interest (*Blm*^*N2*^, *Blm*^Δ*CR1*^, or *Blm*^Δ*CR2*^) balanced over either *TM3, Sb* or *TM6B, Hu Tb* to generate heteroallelic *Blm* females (*e*.*g*., *Blm*^*N1*^ */ Blm*^*N2*^). For experiments with *Blm*^*N1*^ only, *Blm*^*N1*^ *ry e P*{*UASp::Blm*} */ TM6B, Hu Tb* virgin females were crossed *meiP22*^*103*^ *st Blm*^*D2*^ *ry rec*^*1*^ *Ubx P*{*matα::GAL4*} */ TM6B, Hu Tb* males to generate heteroallelic *Blm* null females that could rescue Blm expression after meiotic chromosome segregation to prevent maternal-effect lethality. *Blm*^*D2*^ is another null allele of *Blm* that contains a premature stop codon in the helicase domain (Kusano, 2001). Heteroallelic *Blm* females were then crossed to *y sc cv v g f / Dp(1;Y)B*^*S*^ males. The duplication on the *Y* chromosome carries a dominant mutation causing bar-shaped eyes. Normal progeny resulting from this cross are females whose eyes are *Bar*^*+*^ and males whose eyes are *Bar*^*-*^. Non-disjoined ova that are diplo-*X* result in *XXY* females (and *XXX* progeny who do not survive) whose eyes are *Bar*^*-*^. Non-disjoined ova that are nullo-*X* result in *X0* males (and *Y0* progeny who do not survive) whose eyes are *Bar*^*+*^. *X* NDJ is calculated as the percentage of progeny that arose from NDJ (*Bar*^*-*^ females and *Bar*^*+*^ males), correcting for the loss of half of the diplo- and nullo-*X* ova by multiplying this percentage by two. Crosses were set up as ten females and four males/vial for *Oregon-RM* (wild-type), *Blm*^Δ*CR1*^, and *Blm*^*N2*^ genotypes and thirty females and eight males for *Blm*^Δ*CR2*^ and *Blm*^*N1*^ genotypes. Data were pooled from between 15-60 vials and at least 1000 total progeny to determine the mean NDJ rate for each genotype.

### Mitotic Crossover Assay

Pre-meiotic mitotic crossovers were measured in the male germline using the *net dpp*^*ho*^ *dpy b pr cn* recessive phenotypic marker chromosome. Virgin females with *net dpp*^*ho*^ *dpy b pr cn/ SM6a* and wild-type or various *Blm* alleles (*Blm*^*N2*^, *Blm*^Δ*CR1*^, or *Blm*^Δ*CR2*^) balanced over *TM6B, Hu Tb* were crossed to *w; Blm*^*N1*^*/ TM3, Sb* to generate single males heteroallelic (e.g. *Blm*^*N2*^*/ Blm*^*N1*^) for *Blm* and heterozygous for recessive phenotypic markers for mitotic crossover analysis. For *Blm*^*N1*^ only, virgin females were instead crossed to *w*; *Blm*^*D2*^*/ TM3, Sb*. Single males for each genotype were then crossed to homozygous *net dpp*^*ho*^ *dpy b pr cn* females and scored for mitotic crossovers indicated by the mixed presence and/or absence of recessive phenotypic markers in progeny. Progeny for each single male was scored as a ratio of crossover progeny to total progeny to generate a mitotic crossover rate for each vial. Data for each genotype were pooled from at least 38 vials and 7000 progeny to determine the mean mitotic crossover rate.

### P{w^a^} Assay

The *P*{*w*^*a*^} was performed as described previously (Adams et al., 2003), with minor modifications. First, *y*^*2*^ *w*^Δ^ *P*{*w*^*a*^} virgin females with wild-type or various *Blm* alleles (*Blm*^*N1*^, *Blm*^*N2*^, *Blm*^Δ*CR1*^, or *Blm*^Δ*CR2*^) balanced over *TM6B, Hu Tb* were crossed to *st P*{Δ*2-3*} *Blm*^*D2*^ *Sb/ TM6B, Hu Tb* males to generate single males that were heteroallelic for *Blm* (e.g. *Blm*^*N1*^ */ Blm*^*D2*^) with the *P*{*w*^*a*^} insertion and the Δ2-3 transposase. Single males for each genotype were then crossed to *y*^*2*^ *w*^Δ^ *P*{*w*^*a*^} and progeny were scored for efficiency of repair by resulting eye color: red indicating efficient SDSA, white/yellow indicating end-joining, and apricot indicating no cutting or perfect repair. Progeny from each single male was scored as a ratio of red-eyed progeny to total progeny as a measure of SDSA repair rate. Data for each genotype were pooled from at least 160 vials and 3800 progeny to determine the mean SDSA repair rate.

### Meiotic Crossover Assay

Meiotic crossovers were measured in the female germline using the *net dpp*^*ho*^ *dpy b pr cn* recessive phenotypic marker chromosome. Virgin females with *net dpp*^*ho*^ *dpy b pr cn/ SM6a* and wild-type or various *Blm* alleles (*Blm*^Δ*CR1*^ or *Blm*^Δ*CR2*^) combined with *P*{*matα::GAL4}* for maternal-effect lethality rescue were crossed to *Blm*^*N1*^ *ro e P*{*UASp::Blm*}*/ TM6B, Hu Tb* to generate females heteroallelic for *Blm* and heterozygous for recessive phenotypic markers for meiotic crossover analysis. For *Blm*^*N1*^ only, *net dpp*^*ho*^ *dpy b pr cn/CyO; Blm*^*N1*^ *r e P*{*UASp::Blm*}*/ TM6B, Hu Tb* virgin females were instead crossed to *mei-P22*^*103*^ *st Blm*^*D2*^ *ry rec*^*1*^ *Ubx P*{*matα::GAL4*} */ TM6B, Hu Tb*. Virgin females for each genotype were then crossed to homozygous *net dpp*^*ho*^ *dpy b pr cn* males and scored for meiotic crossovers indicated by the mixed presence and/or absence of recessive phenotypic markers in progeny. Progeny was scored as a ratio of crossover progeny to total progeny to generate a meiotic crossover rate. Data for each genotype were pooled from at least 38 vials and 7000 progeny to determine the mean mitotic crossover rate.

## Acknowledgements

We thank members of the Sekelsky lab for helpful comments on the manuscript. This work was supported by a grant from the National Institute of General Medical Sciences to JS under award 1R35GM118127. EBD was supported in part by a grants from the National Cancer Institute (T32 CA217824). The funders did not play any role in study design, data collection and analysis, decision to publish, or preparation of the manuscript.

The authors declare that they have no competing interests.

## Notes

### Competing Interest Statement

The authors have declared no competing interest.

